# Primary human dermal fibroblasts selectively sense microbial ligands and initiate immune response through chemokines secretion

**DOI:** 10.64898/2026.04.28.721398

**Authors:** Julie Klein, Christophe Gallard, Brigitte David-Watine, Catherine Werts

**Affiliations:** Institut Pasteur, Université Paris Cité, Unité de Biologie et Génétique de la Paroi Bactérienne, F-75015 Paris, France; UTechS Photonic BioImaging, C2RT, Institut Pasteur, Paris, France

**Keywords:** primary human dermal fibroblasts, innate immunity, TLR4/Lipopolysaccharide, Alpk1/ADP heptose, TLR3/Poly I:C, NF-κB activation, IL-8 chemokine

## Abstract

Fibroblasts are traditionally considered structural cells that maintain tissue homeostasis and facilitate repair. However, accumulating evidence suggests they also participate in innate immunity, although their pattern recognition capabilities remain incompletely characterized.

Here, we systematically assessed the innate immune responses of commercially available primary human dermal fibroblasts from a male and a female donor.

Fibroblasts were stimulated with a panel of microbe-associated molecular patterns (MAMPs) targeting various pattern recognition receptors (PRRs), including Toll-like receptors (TLRs), NOD-like receptors (NODs), Alpha kinase 1 (ALPK1) and STING. Innate immune activation was quantified by measuring the nuclear translocation of NF-κB via high content microscopy and cytokines and chemokines secretion by ELISA; baseline PRRs expression was determined by quantitative PCR.

Only a restricted subset of agonists, specifically *E. coli* LPS (TLR4), Poly I:C (TLR3 / RIG-I) and unexpectedly ADP heptose (ALPK1) induced robust NF-κB activation and secretion of the chemokines IL-8 and MCP-1. Apart from IL-6 and RANTES, which were produced exclusively following Poly I:C stimulation, pro-inflammatory cytokines (IL-1β, TNF, IFN-β) and the anti-inflammatory cytokine IL-10 remained undetectable. Consistent with this limited reactivity, qPCR of PRRs revealed basal expression of TLR4 and ALPK1, whereas most other receptors were expressed at very low or undetectable levels. Notably, NOD1 was highly expressed although no cell activation was observed with several NOD1 agonists. Dose-response analysis revealed surprisingly high sensitivity to LPS.

In conclusion, primary human dermal fibroblasts exhibit a highly selective but sensitive innate immune response, largely restricted to chemokine production upon PRR activation. This unexpected dissociation between chemokine and cytokine responses suggests that fibroblasts function as sentinel cells in early skin defense, capable of detecting key microbial patterns at low concentrations, to orchestrate local immune surveillance. Further investigation into interindividual variability and context-dependent activation is needed.

## INTRODUCTION

Fibroblasts are abundant mesenchymal cells that play a central role in extracellular matrix production, tissue remodeling and wound healing (1, 2). However, their role has been re-evaluated in recent years. A growing body of evidence highlights their active participation in immune responses, particularly in the context of chronic inflammation, infection and tissue injury (1, 3–8). Resident cells of most organs, they are strategically positioned to sense changes in their local environment. They interact with immune cells through the release of cytokines and chemokines, suggesting their potential for direct sensing of microbes. Indeed, several studies have shown that fibroblasts express pattern recognition receptors (PRRs), including members of the Toll-like (TLR) and NOD-like receptors (NLR) pathways. Upon activation, these PRRs trigger intracellular signaling cascades that lead to the activation of NF-κB and its translocation from the cytoplasm to the nucleus, where it promotes the transcription of pro-inflammatory cytokines and chemokines. However, the functional relevance of these receptors in fibroblasts remains poorly defined. The literature on this subject is highly heterogenous and presents contradictory results regarding PRR expression, reactivity, and PRR-induced cytokine production. In many cases, conclusions are based on gene expression without functional validation, or on immortalized cell lines, where the immune response may be altered (12), rather than primary human cells. Consequently, the innate immune response of primary fibroblasts remains poorly characterized. To better define the extent of the innate immune response of human fibroblasts, we exposed primary human dermal fibroblasts, from male and female donors to a panel of microbial ligands that target the TLRs, NOD1/2 and ALPK1 pathways. We then quantified their response by measuring nuclear translocation of NF-κB and secretion of cytokines and chemokines.

## MATERIALS AND METHODS

### Ethic statement

All experiments were performed in accordance with relevant guidelines and regulations. The human primary cells used in this study were purchased from TebuBio and are derived from the skin of donors who have given informed consent for the use of their cells in research. Since these cells have been purchased from a certified commercial provider and were anonymized, no further ethical approval was needed.

### Cell culture

Dermal fibroblasts from healthy Caucasian donors used in this study were from TebuBio. The DFM062922A cells (named as DFF-629) and the DFM091119A cells (named as DFF-911) were obtained from the abdomen of a 54-year-old man, and from the arm of a 41-year-old woman, respectively.

Cells were cultured in TPP T75 flasks in 10 mL DMEM GlutaMAX (Gibco) supplemented with 10% decomplemented fetal bovine serum (FBS) (Gibco) (complete medium) at 37°C with 5% CO2. Cells were passaged at 70% confluency. Medium was aspirated and cells rinsed with 5 ml of PBS. Cells were detached with 2 mL of Accutase (StemCell) for 5 min at 37°C 5% CO2, then 3 mL of complete medium was added, and cells were centrifugated for 5 min at 200 g. Cells were resuspended in complete medium and enumerated using a Malassez counting chamber. Viability was estimated using trypan blue. On average, a passage corresponds to a threefold increase in cell population over a period of one week. The cells were routinely analyzed between passages 5–6 and 10. On a regular basis, cells were tested for mycoplasma with the MycoAlert mycoplasma kit (Lonza).

### Fibroblasts stimulation

According to the “Add-only protocol” developed in the laboratory (9), cells were seeded in flat bottom 96-well plates (Greiner 655090) for microscopy analysis or in 96-well plates for cell culture (TPP) at a concentration of 8 000 cells in 100 μL per well the day before the experiment. Agonists used were High molecular weight Poly I:C, lipopolysaccharide from *Escherichia coli* 0111:B4 LPS-EB (tlrl-eblps) and ultrapure (tlr-3pelps), CL429, R848, FLA-ST Ultrapure, ADP-Heptose, c-di-GMP, CpG ODN 2395, CpG ODN 2395 control, Muramyl dipeptide (MDP), MDP control, Murtridap, synthetic diacylated lipopeptide (Pam2CSK4), synthetic tri acylated lipopeptide (Pam3CSK4), all from InvivoGen, whereas β-glucan from *Saccharomyces cerevisiae* was from (Sigma-Aldrich). Concentrations of agonists used and PRR targets are presented in table 1.

**Table 1.** List of the PRRs agonists used with corresponding concentrations and targets.

| **Agonist** | **Used concentration** | **Target** |
| --- | --- | --- |
| **Poly I: C HMW** | 10 µg/mL | TLR3/ RIG-I |
| **LPS E. coli (EB and ultrapure)** | 1 µg/mL<br>Or a range for indicated experiments | TLR4 |
| **CL429** | 1 µg/mL | TLR2 and NOD2 |
| **R848** | 1 µg/mL | TLR7 and TLR8 |
| **Ultrapure flagellin** | 1 µg/mL | TLR5 |
| **ADP heptose** | 1 µg/mL | ALPK1 |
| **C-di-GMP** | 100 µg/mL | STING |
| **CPG ODN 2395** | 15 µg/mL | TLR9 |
| **MDP** | 1 µg/mL | NOD2 |
| **Murtridap**<br>**C12-iE-DAP** | 250 nM<br>250 nM, 375 nM, 500 nM | Human NOD1 |
| **Pam2CSK4** | 1 µg/mL | TLR2/TLR6 |
| **Pam3CSK4** | 1 µg/mL | TLR2/TLR1 |
| **β-glucan** | 1 µg/mL | Dectin-1 |

For stimulation, agonists were diluted in 100µl DMEM GlutaMAX supplemented with 10% decomplemented FBS at twice the final stimulation concentration. Stimulation of fibroblasts was performed by adding 100 μL of stimulation reagent per well and incubated at 37°C in 5% CO2 for 1 h (for indicated experiments) or 3 h. Each stimulation was performed in triplicate and repeated at least three times in independent experiments for each male and female fibroblast cells. For cytokines analysis, supernatants were collected 24 h post-infection, transferred in a flat bottom 96-well plate and kept at -20°C. For high-content microscopy processing, after 1 h (for indicated cells) or 3 h stimulation, cells were fixed with paraformaldehyde (PFA) directly added to the medium to a final concentration of 4% for 10 min. The fixator was removed by aspiration, and cells were kept at 4°C in PBS.

### Immunofluorescence labelling, high content microscopy and analysis

Cells were permeabilized with PBS 1X-Triton 0,2% D-PBS (Life Technology) for 15 min. After removal of the permeabilization buffer, cells were incubated in blocking buffer (PBS 1X-fat dry milk 3,5%) for 1 h at room temperature under agitation and incubated overnight at 4°C with a monoclonal anti-human NF-κB p65 antibody (27F9.G4, Rockland) diluted 1: 1000 in PBS 1X-fat dry milk 3,5%. Cells were then stained for 2 h at room temperature under agitation with goat anti-mouse Alexa-555 coupled secondary antibody (A-21422, Invitrogen) diluted 1:500 in PBS 1X-fat dry milk 3,5%. Finally, nuclei were stained with Hoechst 33342 (Invitrogen) for 20 min at room temperature under agitation (0,5 µg/mL in PBS 1X) before being washed with PBS 1X for the microscopy analysis.

Images were acquired on the automated spinning disk confocal microscope Opera Phenix (Perkin Elmer Technologies, UtechS PBI, Institut Pasteur) in non-confocal mode. Images of 13 fields from 3 independent wells per condition (roughly covering the entire surface of each well to obtain a reliable statistical analysis) were acquired at 10x magnification air with a NA= 0,3. The fluorophores were detected using the following settings: Hoechst 33342 (excitation wavelength 405 nm; filter 450/50), and Alexa 555 (excitation wavelength 561 nm; filter 600/40). Images analysis was performed using Signal Image Artist (Perkin Elmer), with the following steps: (i) input image; (ii) find nuclei using Hoechst 33342 signal; (iii) find cytoplasm with Alexa 568; (iv) select region, nuclei are resized (20%); (v) calculate the area and the roundness of the resized nuclei; (vi) calculate the area of the cytoplasm; (vii) select population (120 µm2 > nucleus area < 450 µm2; nucleus roundness 0,82; cytoplasm area < 4500 µm^2^); (viii) calculate cytoplasm intensity; (ix) calculate nuclei intensity; (x) calculate nuclei intensity/ cytoplasm intensity ratio; (xi) formulate calculated outputs : % positive cells (ratio>1,2) ; (xii) formulate calculated outputs : % negative cells (ratio<1,2). To determine this threshold, preliminary experiments quantified NF-κB nucleus to cytoplasm intensity ratios at the single-cell level using Columbus software (PerkinElmer) and statistically analyzed with GraphPad Prism, with outliers removed using the ROUT method (Q=1%). A total of 63 872 cells were analyzed across three conditions (control: 21 761 cells; LPS: 20 761 cells; Poly I:C: 21 350 cells) following a 3-hour stimulation. Kruskal-Wallis multiple comparison testing revealed significant differences between control and treated populations (p< 0.0001). Control cells exhibited ratios ranging from 0.8 to 1.2, with cumulative distributions showing that 99% of cells remained below a ratio of 1.2, whereas LPS and Poly I:C treated cells predominantly displayed ratios exceeding 1.2. Consequently, a threshold of 1.2 was established to define cellular activation status, with cells having ratios ≥1.2 classified as activated and those <1.2 as non-activated.

### ELISA

The quantification of human IL8, RANTES, IL-6, MCP-1, IL1β, IL-10, TNF and IFN-β released in cell supernatants was performed using ELISA DuoSet kits (R&D Systems) according to the manufacturer’s instructions. Supernatants were kept at -20°C before cytokine dosage. Samples were tested at least in triplicate and standards were included in every plate.

### Quantification of gene expression by RT-qPCR

Total RNAs were prepared from 2,5 × 10^!^ cells and purified with the RNeasy Mini Kit (Qiagen) according to the supplier’s instructions. RNA concentration and purity were assessed with a NanoDrop 2000 spectrophotometer (Thermo Fisher Scientific) at OD260nm. Then, their concentrations were adjusted to 0,1 µg/µL with RNAse free water and then kept at -80°C.

For cDNA synthesis, 1 µg of RNA was transcribed using the SuperScript II reverse transcriptase according to the manufacturer’s instructions (Invitrogen). cDNAs were stored at -20°C. Quantitative RT-PCR was performed on a StepOne Plus real-time PCR machine (Applied Biosystems). 0,2 ng of cDNA were distributed in duplicate and mixed with the TaqMan Universal Master Mix II (Applied Biosystem) with primers and probes, which are listed in Table 2. Ct values were normalized by subtracting the HPRT Ct in each sample.

**Table 2.** Targets and probes list used for quantitative RT-PCR.

| Target | Probe reference |
| --- | --- |
| HPRT | Pre-designed: **Hs02800695_m1** (ThermoFisher) |
| TLR1 | Pre-designed: **Hs00413978_m1** (ThermoFisher) |
| TLR2 | Pre-designed: **Hs00610101_m1** (ThermoFisher) |
| TLR3 | Pre-designed: **Hs01551079_g1** (ThermoFisher) (1)<br>Pre-designed: **Hs00152933_m1**(ThermoFisher) (2) |
| TLR4 | Pre-designed: **Hs00152939_m1** (ThermoFisher) |
| TLR5 | Pre-designed: **Hs01019558_m1** (ThermoFisher) |
| TLR6 | Pre-designed: **Hs04975839_m1** (ThermoFisher) |
| CLEC7a | Pre-designed: **Hs00224028_m1** (ThermoFisher) |
| CD14 | Pre-designed: **Hs001669122_g1** (ThermoFisher) |
| CD36 | Pre-designed: **Hs00354519_m1** (ThermoFisher) |
| ALPK1 | Pre-designed: **Hs01567926_m1** (ThermoFisher) |
| MD2 | Pre-designed: **Hs00209770_m1** (ThermoFisher) |
| NOD1 | Pre-designed: **Hs01036720_m1** (ThermoFisher) |
| NOD2 | Pre-designed: **Hs01550753_m1** (ThermoFisher) |
| RIG-I | Pre-designed: **Hs01061436_m1** (ThermoFisher) |

### Statistical analysis

All statistical analyses were performed using GraphPad Prism (version 10.0, GraphPad Software, San Diego, CA). For comparisons between stimulated and non-stimulated (control) conditions, a one-way analysis of variance (ANOVA) was conducted. Each experimental condition was compared individually to the non-stimulated control using Dunnett’s multiple comparisons post hoc test. A p-value of less than 0,05 was considered statistically significant. For specific pairwise comparisons, un unpaired two-tailed Student’s t-test was performed.

## RESULTS

### Primary human fibroblasts activate NF-κB

We have previously demonstrated that NF-κB translocation is a robust read-out for monitoring pattern recognition receptor (PRR) responses in primary human dermal fibroblasts from healthy donors of the LabEx Milieu Interieur cohort (9). In this context, NF-κB activation was assessed at 3 hours after stimulation (hps), a different timeframe from the faster kinetics typically observed in other cell types such as human macrophages, where activation reaches its maximum around 1 hps (10) (11). To determine whether this delayed kinetic profile could be observed in the primary fibroblasts used in this study (DFF-911 and DFF-629 cells), cells were stimulated with *E. coli* LPS or Poly I:C and analyzed after 1h and 3h by high-content microscopy (Fig 1A). Representative microscopy images illustrate the nuclear translocation of NF-κB after stimulation (Fig 1B). While non-stimulated fibroblasts display a predominant cytoplasmic localization of NF-κB, LPS-stimulated cells exhibit a marked enrichment of the NF-κB signal in the nucleus confirming the validity of this read-out at the single cell level. The percentages of positive cells with NF-κB in the nucleus were quantified. NF-κB nuclear translocation was detected after stimulation in both conditions, with no statistically significant difference between 1 h and 3 h (Fig 1A) Despite the lack of significant difference, we maintained a 3 h duration for subsequent experiments to ensure consistency with our previous work on primary human fibroblasts (9). As a control, we performed the same experiment in L929 murine dermal fibroblast cell line, which showed a classical NF-κB peak of activation at 1 hps and not at 3 hps (data sup 1). This could reflect specific cell type and host-related differences in LPS sensing.

**Fig 1.**
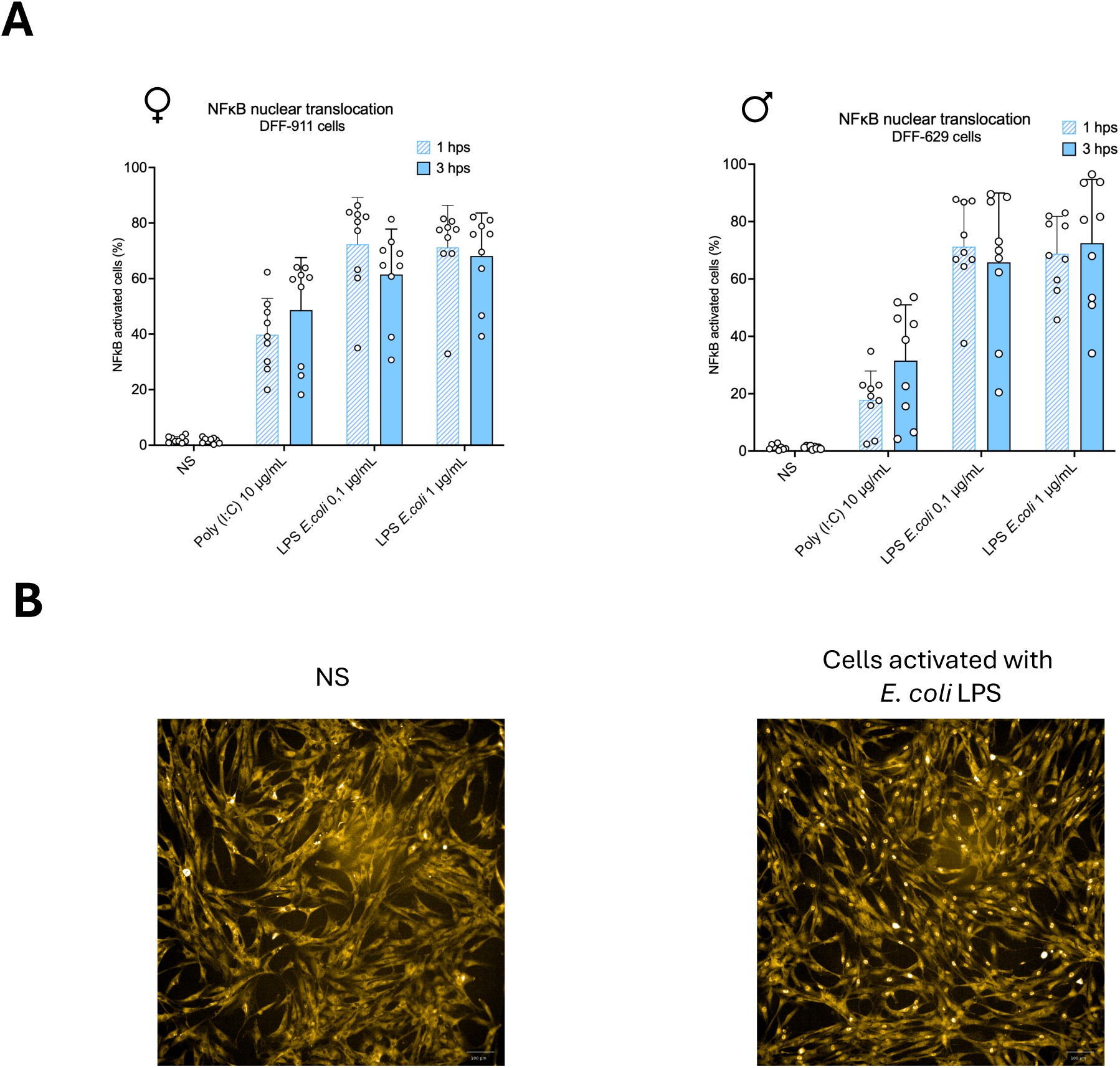
NF-kB nuclear translocation in primary human dermal fibroblasts. Primary Human dermal fibroblasts were stimulated with *E. coli* LPS or Poly I:C for 1 hour and 3 hours. Cells were labeled with an anti NF-κB antibody and images were acquired at 10X magnification. (A) Quantification of the percentage of cells displaying nuclear NF-κB (activated). Data from three independent experiments were pooled; each experimental condition includes three technical replicates per experiment. Bars represent mean ± SD of the biological replicates. Statistical analysis was performed using t-test between each condition. Non-significant comparisons are not indicated. (B) Representative fluorescence microscopy images of non-stimulated (NS) and LPS *E. coli* stimulated cells 3 hps. Scale bar, 100 µm.

### Only *E. coli* LPS, Poly I: C and ADP heptose induce NF-κB activation, IL8 and MCP-1 secretion

To determine the innate immune responsiveness of primary human dermal fibroblasts, cells from two independent donors, a woman (DFF-911) and a man (DFF-629) were stimulated with a panel of agonists specifically targeting different pattern recognition receptors (PRRs). These included ligands for TLR2/1 (Pam3CSK4), TLR2/6 (Pam2CSK4), TLR3 /RIG-I(Poly I :C), TLR4 (LPS from *E.coli*), TLR5 (ultrapure flagellin), TLR7/8 (R848), TLR9 (ODN2395 and control), NOD1 (M-TriDaP), NOD2 (MDP and control), TLR2/NOD2 (CL429), ALPK1 (ADP heptose), STING (cyclic di-GMP) and Dectin-1 (β-glucan). NF-κB activation was measured 3 hps by high-content microscopy and chemokines / cytokines secretion by ELISA.

In both donors, only *E. coli* LPS, Poly I: C and ADP-heptose induced a clear and reproducible NF-κB nuclear translocation (Fig 2A and data sup 2), as well as strong secretion of IL-8 and MCP-1 (Fig 2B and 2C). β-glucan induced a very weak NF-κB signal, but this response was marginal and not accompanied by IL-8 secretion. All other agonists failed to elicit NF-κB activation or cytokine production.

**Fig 2.**
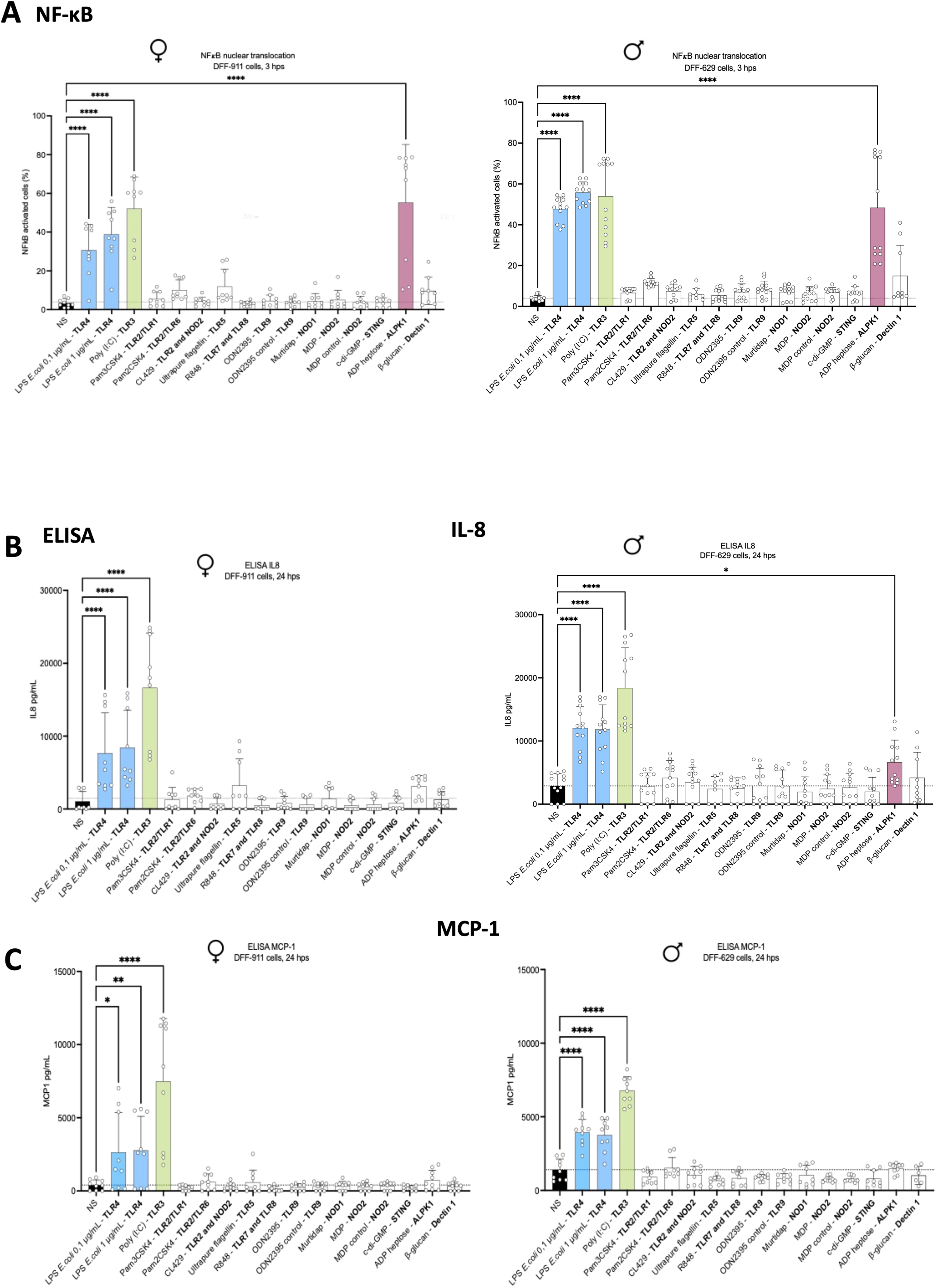
Response of primary human fibroblasts to a panel of PRR agonists. Primary human dermal fibroblasts from two independent donors (DFF-911, female and DFF-629, male) were stimulated by a panel of PRR agonists for 3 hours. (A) Cells were labeled with an NF-κB antibody and images were acquired with a 10X magnification and the NF-κB translocation was quantified (n=3). (B) IL-8 and (C) MCP-1 production was measured by ELISA 24 hps. Experiment representative of n=3 (DFF-911) and n=4 (DFF-629) with three biological replicates per condition. Bars represent mean ± SD of the biological replicates. Statistical analysis was performed using one-way ANOVA followed by Dunnett’s multiple comparisons test against the non-stimulated control. Only statistically differences are indicated (p < 0,05); non-significant comparisons are not marked.

Unexpectedly, IL-8 and MCP-1 were the only chemokine consistently produced in both cell types. IL-6 and RANTES were secreted only by Poly I:C, while IL-1β, TNF-⍺, IFN-β and IL-10 were undetectable under all conditions (data sup 3).

These results highlight a highly selective innate immune response profile in primary human dermal fibroblasts. This response is characterized by activation of the NF-κB pathway and the production of IL-8 and MCP-1 in response to a limited number of stimuli. The responses of male and female donors are very similar, suggesting that this restricted sensing phenotype is a common feature of human dermal fibroblasts.

### qPCR reveals a restricted basal expression pattern consistent with the narrow functional responsiveness of fibroblasts

To determine whether the selective NF-κB activation observed in response to LPS, Poly I:C and ADP-heptose reflected an underlying transcriptional landscape, basal expression of key PRR genes was quantified in unstimulated fibroblasts from both donors (DFF-911 and DFF-629). RT-qPCR analysis revealed a restricted transcriptional profile in the two donors (figure 3). Both fibroblasts expressed *alpk*1, which is consistent with the recognition of ADP-heptose, as well as *cd14* and *md2*, in agreement with the recognition of LPS, although the expression of *tlr4* was minimal. The cytosolic receptor *nod1* was highly expressed, although we did not observe response to the NOD1 agonist, Mur tri-DAP (Fig 2A and 2B), neither to the soluble NOD1 agonist C12-iE-DAP (sup Fig 4). *tlr6* was also expressed, unlike the genes encoding CD36, TLR2 and TLR1. These results are consistent with the lack of recognition of the Pam3cys agonist via TLR2 and TLR1, and the very weak response observed with Pam2cys, a TLR2/6 agonist often associated with CD36 (12). *CLEC7A* (Dectin-1), *tlr5* (Flagellin), *tlr9, sting* and *tlr3* were either not detected or were detected only at a very low level. The low expression of *tlr3* was particularly surprising, given the strong response of the cells to Poly I: C. To determine whether *tlr3* expression could be activated only after stimulation, we measured its expression in cells stimulated for three hours with Poly I: C. However, despite using two different primer sets for *tlr3*, we did not detect *tlr3* expression (Sup Fig 4), suggesting a RIG-I response, a cytosolic receptor involved in Poly I:C recognition. Expression of the RIG-I gene was therefore tested. The RIG-I gene was expressed in cells from both donors (sup Fig 5).

**Fig 3.**
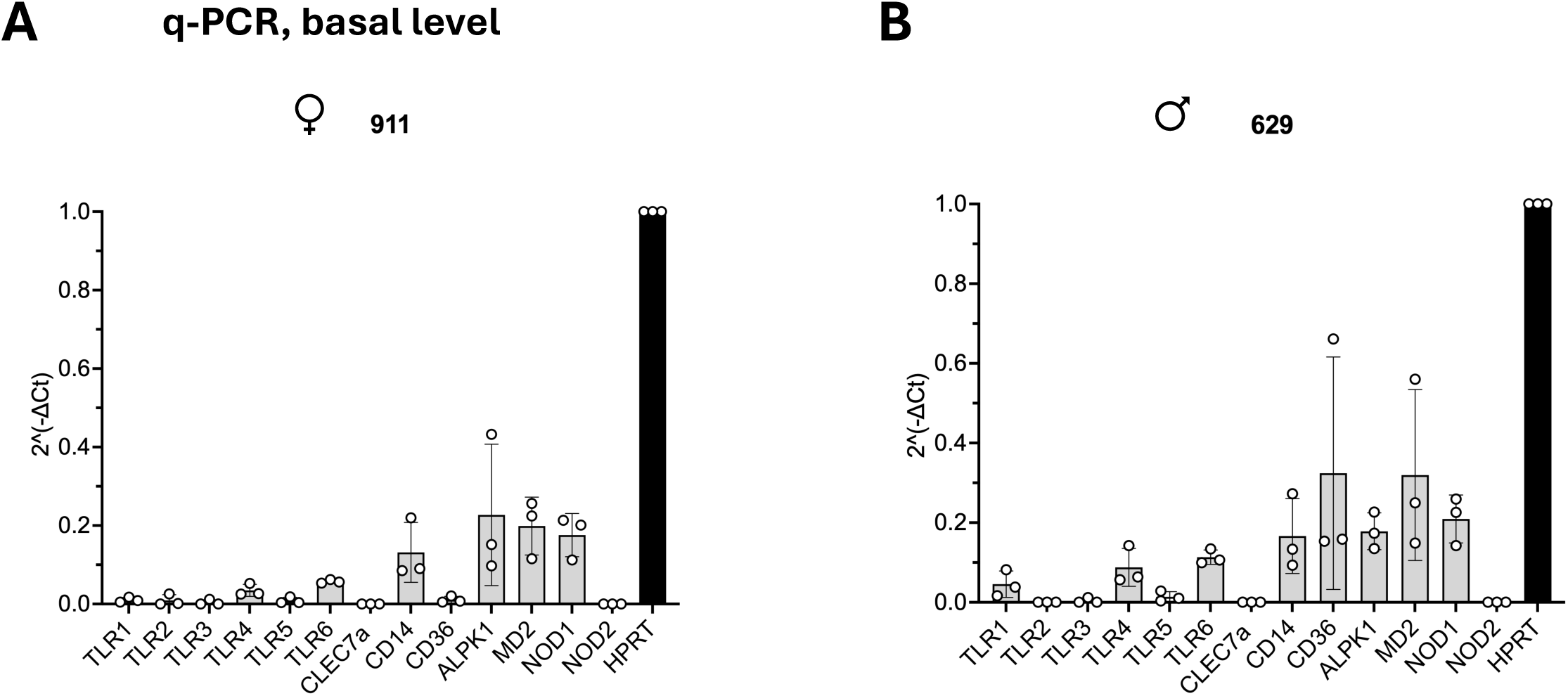
PCR analysis of PRR expression in human dermal fibroblasts at a basal level. Basal expression of PRRs and associated co-receptors in unstimulated primary human dermal fibroblasts. mRNA levels of TLR1, TLR2, TLR3, TLR4, TLR5, TLR6, CLEC7A, CD14, CD36, ALPK1, MD2, NOD1 and NOD2 were quantified by RT-qPCR in fibroblasts from a female donor (DFF-911) and a male donor (DFF-629). Expression is shown as 2^(-ΔCt) relative to HPRT (set to 1). Each data point represents one fully independent biological replicate, consisting of an independent cell culture, RNA extraction, cDNA synthesis and PCR reaction (n=3). Bars represent mean ± SD.

### Primary human fibroblasts are highly sensitive to low doses of *E. coli* LPS

Since we did not observe a dose response effect to LPS used at 100 ng compared to 1 µg/mL (Fig 2), we sought to determine the sensitivity threshold of primary human dermal fibroblasts to *E. coli* LPS. Therefore, we stimulated cells from the two donors (DFF-629, male and DFF-911, female) with decreasing LPS concentrations, ranging from 10 ng/mL to 0,01 ng/mL. Two LPS preparations were used: standard EB LPS and ultrapure LPS, both derived from *E. coli*. Nuclear translocation of NF-κB was measured 3 hps by high content microscopy. Both DFF-629 and DFF-911 cells responded to LPS in a dose-dependent manner, with a NF-κB activation observed at 1 ng/mL for both LPS preparations (Fig 4A et 4B). Below this threshold, NF-κB activation decreased significantly. In the male donor (DFF-629), NF-κB activation was undetectable at 0,1 ng/mL and 0,01 ng/mL. In contrast, fibroblasts from the female donor (DFF-911) retained a measurable, albeit not significant, NF-κB response at 0,1 ng/mL, indicating slightly higher sensitivity. In both donors, standard EB LPS elicited a stronger NF-κB response than the ultrapure form at all concentrations of *E. coli* LPS. This difference likely reflects the presence of additional immune-stimulatory contaminants in the standard preparation (13). These findings demonstrate that primary human dermal fibroblasts are highly sensitive to low concentrations of *E. coli* LPS and that donor-related factors as well as the purity of the LPS preparation, influence the sensitivity and activation threshold of NF-κB.

**Fig 4.**
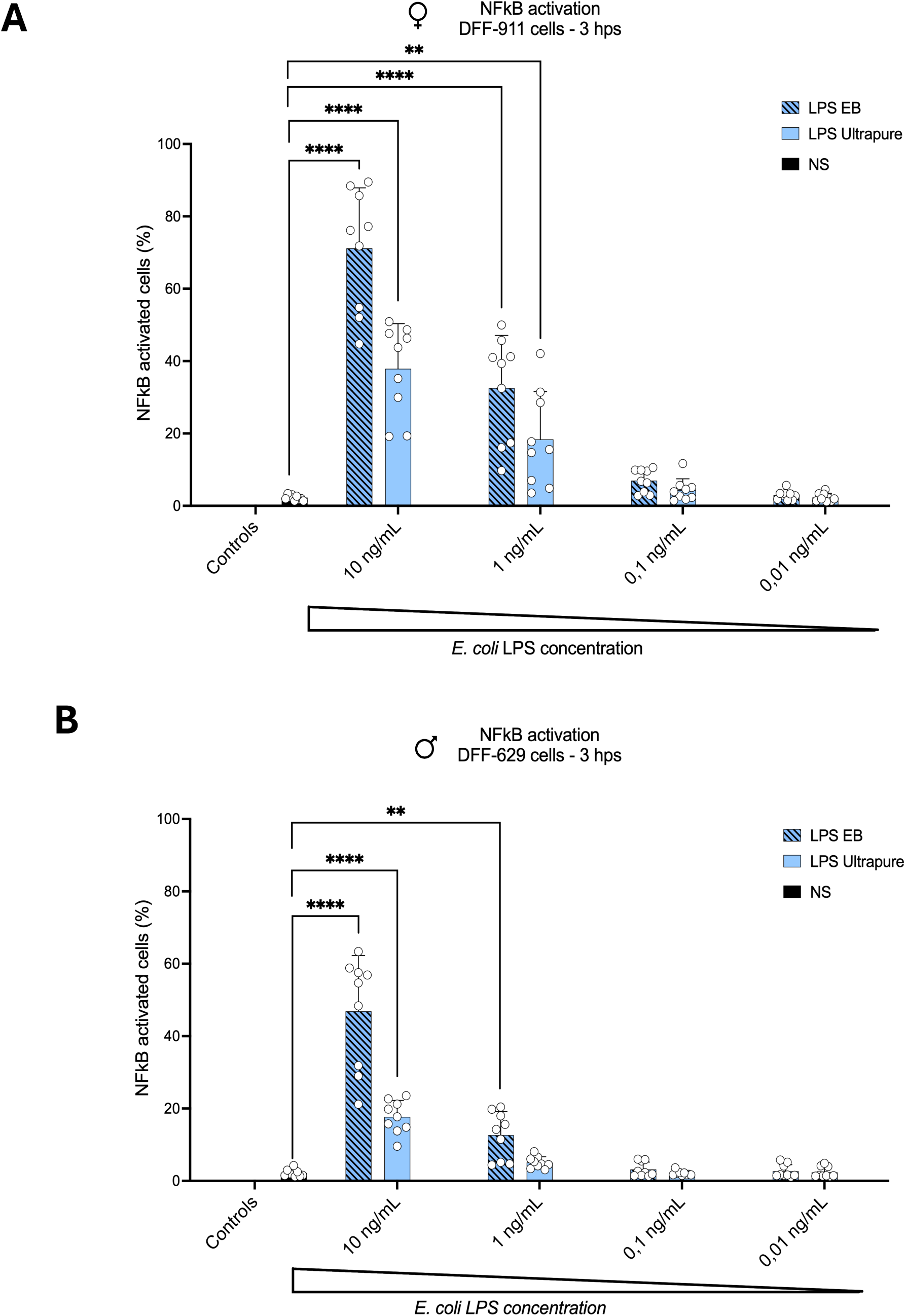
NF-κB nuclear translocation in response to standard EB and ultrapure LPS. Primary human dermal fibroblasts from two independent donors (DFF-911, female and DFF-629, male) were stimulated for 3 hours by two types of *E. coli* LPS, the standard LPS EB and the ultrapure form. NF-κB translocation was quantified in (A) DFF-911 cells and in (B) DFF-629 cells. Experiment representative of n=3 with three biological replicates per condition. Error bars correspond to the standard deviation of three biological replicates. Statistical analysis was performed using one-way ANOVA followed by Dunnett’s multiple comparisons test against the non-stimulated control. Only statistically differences are indicated (p < 0,05); non-significant comparisons are not indicated.

In conclusion, our results revealed a surprisingly selective yet highly sensitive innate immune activation profile in primary human dermal fibroblasts. Responses were limited to RIG-I, TLR4 and ALPK1 signaling pathways, mainly inducing the production of IL-8 and MCP-1 chemokines. Furthermore, we have shown that these fibroblasts were highly sensitive to *E. coli* LPS responding to concentrations as low as 1 ng/mL.

## DISCUSSION

In this study, we characterized the innate immune response of primary human dermal fibroblasts, focusing on the NF-κB activation pathway and the production of the chemokines IL-8 and MCP-1. Our results reveal that these cells display a highly selective response to microbial stimuli, with robust activation observed in response to *E. coli* LPS, Poly I:C, and ADP-heptose. This activation was consistently associated with the secretion of IL-8 and MCP-1, highlighting a restricted but functionally coherent inflammatory program. We also demonstrated their remarkable sensitivity to LPS, comparable to that of monocytes (14, 15).

This selective response suggests that dermal fibroblasts possess a limited repertoire of functional PRRs. Indeed, most the agonists tested, including ligands of TLR2, TLR5, TLR7/8, TLR9, NOD1/NOD2 and STING, did not induce NF-κB activation or cytokine production. This observation is corroborated by our RT-qPCR data showing low, or even undetectable expression of many corresponding receptors. This restricted detection profile indicates that fibroblasts, unlike professional immune cells, exhibit a limited PRR expression pattern, likely reflecting their specialized role in tissue homeostasis rather than large scale pathogen detection (16).

Among the signaling pathways involved, ADP-heptose induced activation deserves special attention. This bacterial metabolite is detected via the ALPK1-TIFA-NF-κB axis. Although this pathway is well described in epithelial cells, particularly in the context of *Helicobacter pylori* infection (17), its functionality in primary human dermal fibroblasts remains poorly documented. Our results provide evidence that this pathway is active in dermal fibroblasts, suggesting that these cells can detect intracellular bacterial metabolites and contribute to early inflammatory signaling in infected tissues. The mechanism by which extracellular ADP-heptose reaches the fibroblast cytosol remains unclear but may involve passive uptake, membrane perturbation, or another -yet-unknown passive mechanism.

A striking observation from our study is the discrepancy between receptor expression and functional response for certain signaling pathways, notably TLR3. Despite clear NF-κB activation in response to Poly I:C, TLR3 transcripts were undetectable or present at extremely low levels, even with different primer sets. This suggests that Poly I:C sensing in these fibroblasts may be independent of TLR3. In this context, the expression of RIG-I, a cytosolic RNA sensor capable of recognizing double-stranded RNA, represents a plausible alternative pathway. Our results support the hypothesis that RIG-I could play a major role in Poly I:C sensing by dermal fibroblasts.

However, the role of TLR3 in the response to Poly I:C in fibroblasts has been demonstrated in human fibroblasts, notably by the study of human cells carrying mutants in this pathway, and in the *tlr3* gene (9, 12). It also remains possible that very low levels of TLR3 expression are sufficient for signaling.

Another notable finding concerns NOD1. Although NOD1 transcripts were highly expressed in both donors, stimulation with specific agonists failed to induce NF-κB activation or cytokine production. This suggests that NOD1 signaling is not functionally active under our experimental conditions. There are several possible explanations for these observations. Firstly, NOD1 activation requires the intracellular delivery of its ligand; inefficient uptake of Mur-Tri dap could therefore limit signaling. However, using C12-iedap, a soluble, fatty version of the NOD1 agonist ieDAP, did not either activate the fibroblasts. Second, adaptor or signaling molecules essential downstream of NOD1, such as RIPK2, could be under expressed or inactive in these cells. Finally, regulatory mechanisms could actively suppress NOD1 signaling in fibroblasts to prevent inappropriate activation. This observation underscores the importance of distinguishing gene expression from functional signaling capacity when assessing innate immune pathways.

Among classical cytokines expressed after PRR stimulation, and tested in this study, IL-8 and MCP-1 chemokines were predominantly produced after stimulation. IL-8 is a key chemoattractant for neutrophils, while MCP-1 promotes monocytes recruitment, indicating that fibroblasts may contribute to the early stages of immune cell infiltration at site of infection or tissue injury. The limited production of other cytokines, such as IL-1β, TNF-⍺, IFN-β or IL-10, supports the hypothesis of a finely regulated inflammatory response. This restricted cytokine profile suggests that fibroblasts are specialized in recruiting immune cells rather than orchestrating a generalized inflammatory response. The selective induction of IL-6 and RANTES solely by Poly I:C suggests that antiviral-like responses may engage additional transcriptional programs not activated by bacterial stimuli.

Finally, our data demonstrate that primary human dermal fibroblasts exhibit remarkable sensitivity to *E. coli* LPS, with detectable NF-κB activation at concentrations as low as 1 ng/mL in both donors. Interestingly, (the female) donor fibroblasts (DFF-911) showed a modest NF-κB response even at 0,01 ng/mL, suggesting some interindividual variability in the response to LPS as described in Chansard et al. This sensitivity is comparable to that described for classical innate immune cells such as monocytes and macrophages (10) indicating that fibroblasts, traditionally considered as structural cells, may play a more active role in detecting bacterial components in tissues. Their reactivity to very low doses of LPS makes them important sentinels capable of detecting bacterial invasion or tissue damage. The more intense response elicited by the standard EB LPS preparation compared to that obtained with ultrapure LPS, suggests that contaminating molecules, such as lipoproteins or other TLR2 agonists, could act in synergy with the TLR4 signaling pathway to amplify the NF-κB response (13). This observation underscores the importance of LPS purity in the design and interpretation of experiments, particularly when studying the response at low doses (18). The variability observed between donors, although limited, is consistent with the growing body of evidence indicating that genetic and epigenetic differences can influence the expression of innate immune receptors and the signaling threshold. Although fibroblasts are not traditionally considered as innate immune cells, they can nevertheless contribute to *in situ* inflammatory responses by detecting microbial products.

Beyond their role in tissue structure and repair, our results confirm the hypothesis that dermal fibroblasts can act as functional immune sentinels, particularly during the early stages of skin infection. In situations such as open wounds contaminated with environmental pathogens, fibroblasts are among the first cells to meet microbial components in the dermis. This strategic location allows them to detect microbial invasion and rapidly initiate an inflammatory response.

Together, our results provide a detailed characterization of the innate immune response capacity of primary human dermal fibroblasts and highlight the importance of considering these cells as key players in host defense and infection response. Their ability to selectively sense bacterial products and secrete chemokines such as IL-8 and MCP-1 positions them as essential initiators of neutrophil and monocyte recruitment. In a tissue environment like the skin, where early pathogen containment is crucial the role of fibroblasts signals could attract the classical immune cells to reach the infected area. By defining the specificity and sensitivity of PRR mediated signaling in fibroblasts, this study provides a clearer framework for understanding their immunological role in skin infection.

### Limitations and perspectives

This study presents several limitations. First, it relies on fibroblasts isolated from only two donors, which, despite consistent trends, limits the generalizability of the results given the known heterogeneity of innate immune response among individuals (9, 19). Furthermore, although we tested a wide range of PRRs agonists, it is possible that other relevant stimuli, alone or in combination could elicit different responses. Moreover, the existing literature on the innate immune functions of fibroblasts is highly heterogeneous, with widely varying results ranging in terms of receptor expression and functional reactivity, depending on tissues origin, culture conditions, and interindividual variability. This diversity complicates direct comparison of studies and underscores the need for standardized approaches.

Future research should include a larger donor cohort, explore additional functional parameters such as transcriptional profiling or secretome analysis, and consider experiments on *in vivo* or 3D tissue models to better understand the role of fibroblasts in their microenvironment.

Furthermore, future work should explore the molecular mechanisms underlying the selective response of these cells, including the intracellular localization and the regulation of TLRs pathways. Additional studies using transfection to administer ALPK1 inhibitors would be necessary to confirm the involvement of this cytosolic sensing pathway. Understanding the precise conditions under which fibroblasts participate in the immune response will be essential for harnessing their potential as therapeutic targets or modulators in infectious and inflammatory diseases.

## Acknowledgments

The authors thank Dr Anne Danckaert (biostatistician, UtechS PBI, C2RT, Institut Pasteur) for her expert assistance with the statistical validation of our image analysis method. We also thank all members of the UTechS Photonics Bioelectronic Imaging (PBI) group for their support, technical assistance and advice, and especially Dr. Nassim Mahtal for his help in managing data on Columbus and SIMA as well as for his advice on the optimal use of the Opera-Phenix microscope.

## Author contributions

JK: investigation, data analysis and interpretation, figures, methodology, writing original draft-review editing. CG: investigation, methodology. BDW: investigation, data analysis and interpretation, supervision, writing draft-review editing. CW: conception, investigation, data analysis and interpretation, supervision, visualization, writing draft-review editing.

## Statements

All data are available in the main text or in the Supplementary Material. Raw data and analyses are secured on elabnotebook at Institut Pasteur and will be made available on request to Catherine Werts.

## Declaration of conflicting interest

The authors declare no conflicts of interest in this study.

## Funding statement

This study was supported by the international research coordination research on animal disease (ICRAD) grant S-CR23012-ANR 22 ICRD 0004 01 to CW.

The funders had no role in study design, data collection, data analyses, interpretation, or writing of this article

---

**Data sup 1.**
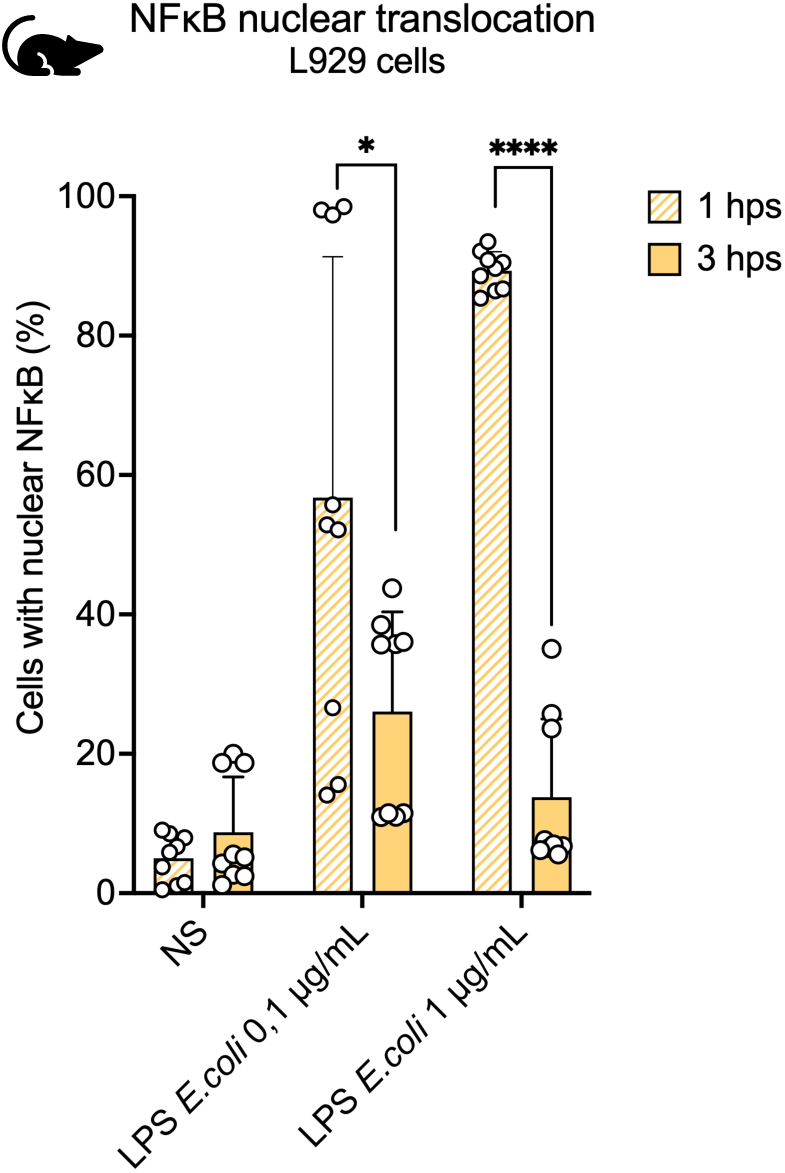
NF-κB kinetics of murine dermal fibroblasts after *E. coli* LPS stimulation. Murine fibroblasts cell line (L929 cells) were stimulated by *E. coli* LPS for 1 hour and 3 hours. Cells were labeled with an anti NF-κB antibody and images were acquired with a 10X magnification. Experiment representative of n=3 with three biological replicates per condition. Error bars correspond to the standard deviation of three biological replicates. Statistical analysis was performed using t-test between each condition. Non-significant comparisons are not marked.

**Data sup 2.**
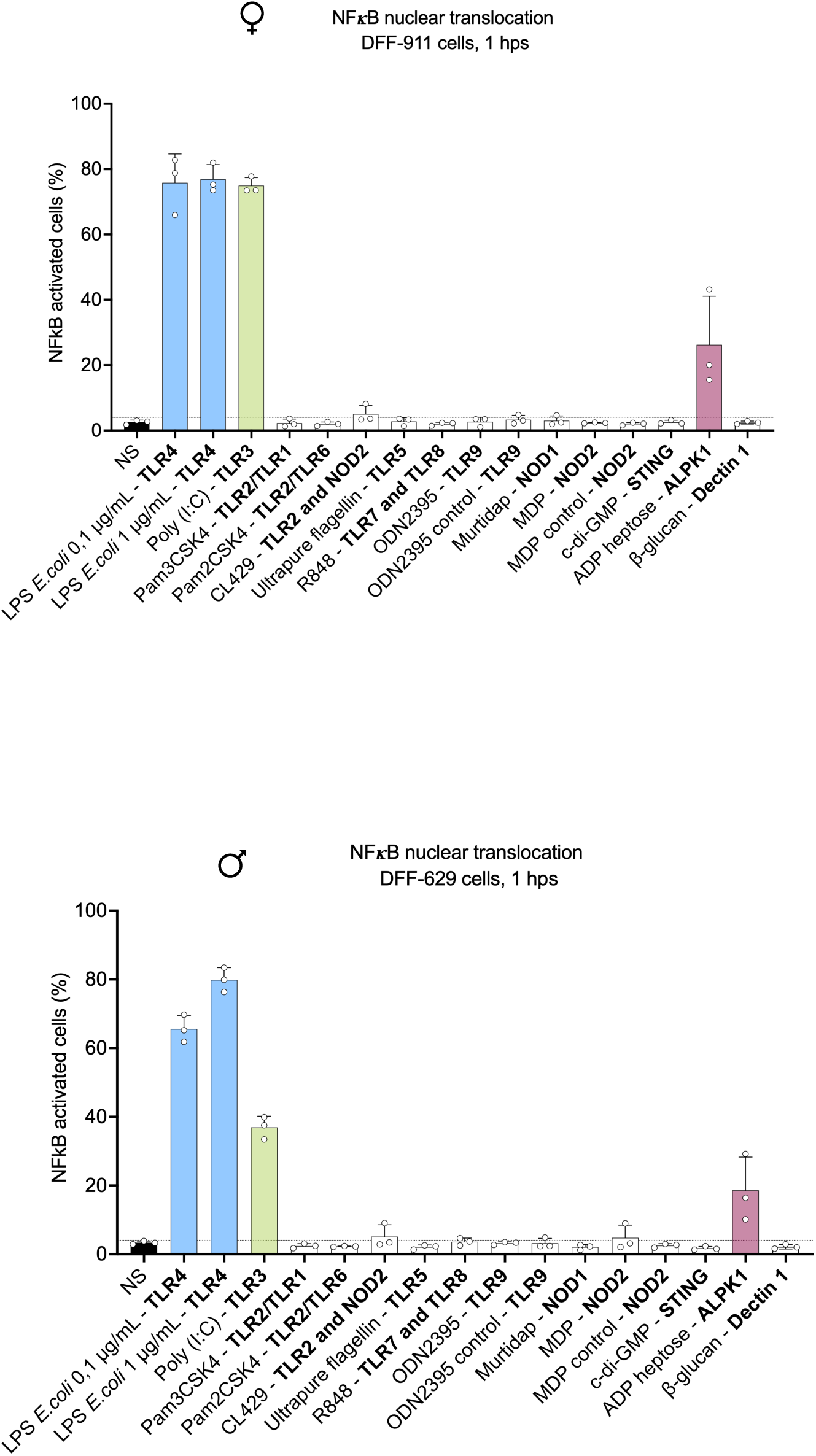
Response of primary human fibroblasts to a panel of PRR agonists 1h post stimulation. Primary human dermal fibroblasts (DFF-911, female and DFF-629, male) were stimulated by a panel of PRR agonists for 1 hour. Cells were labeled with an NF-κB antibody and images were acquired with a 10X magnification and the NF-κB translocation was quantified (n=1 with 3 biological replicates per condition).

**Data sup 3.**
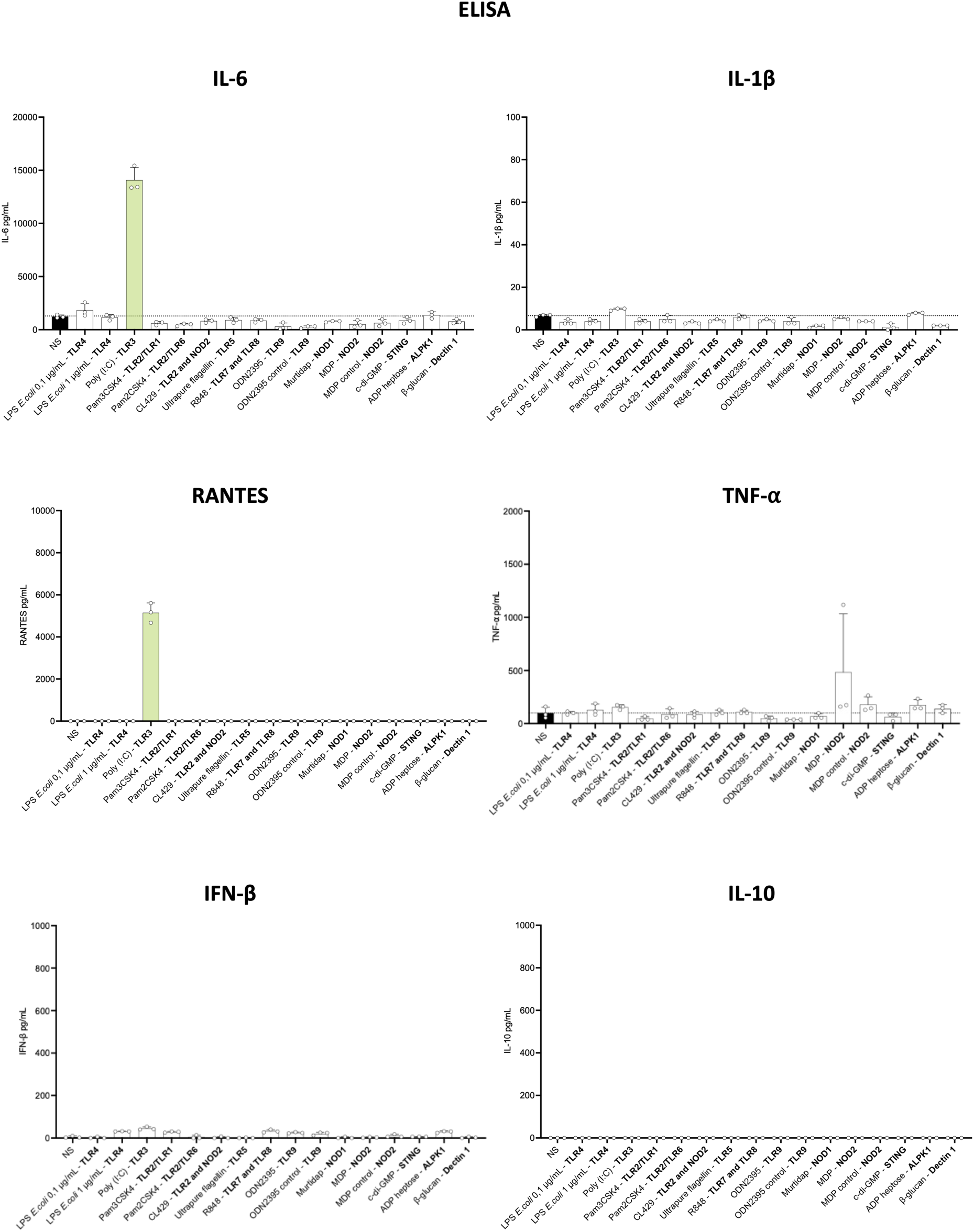
Inflammatory response of human dermal fibroblasts to PRRs agonists. Primary human dermal fibroblasts from a female donor DFF-911 were stimulated with a panel of PRR agonists for 3 hours. IL-6, IL-1β, RANTES, TNF-⍺, IFN-β and IL-10 production was measured by ELISA 24 hps. N=1 with three biological replicates per condition. Bars represent mean ± SD of the biological replicates.

**Data sup 4.**
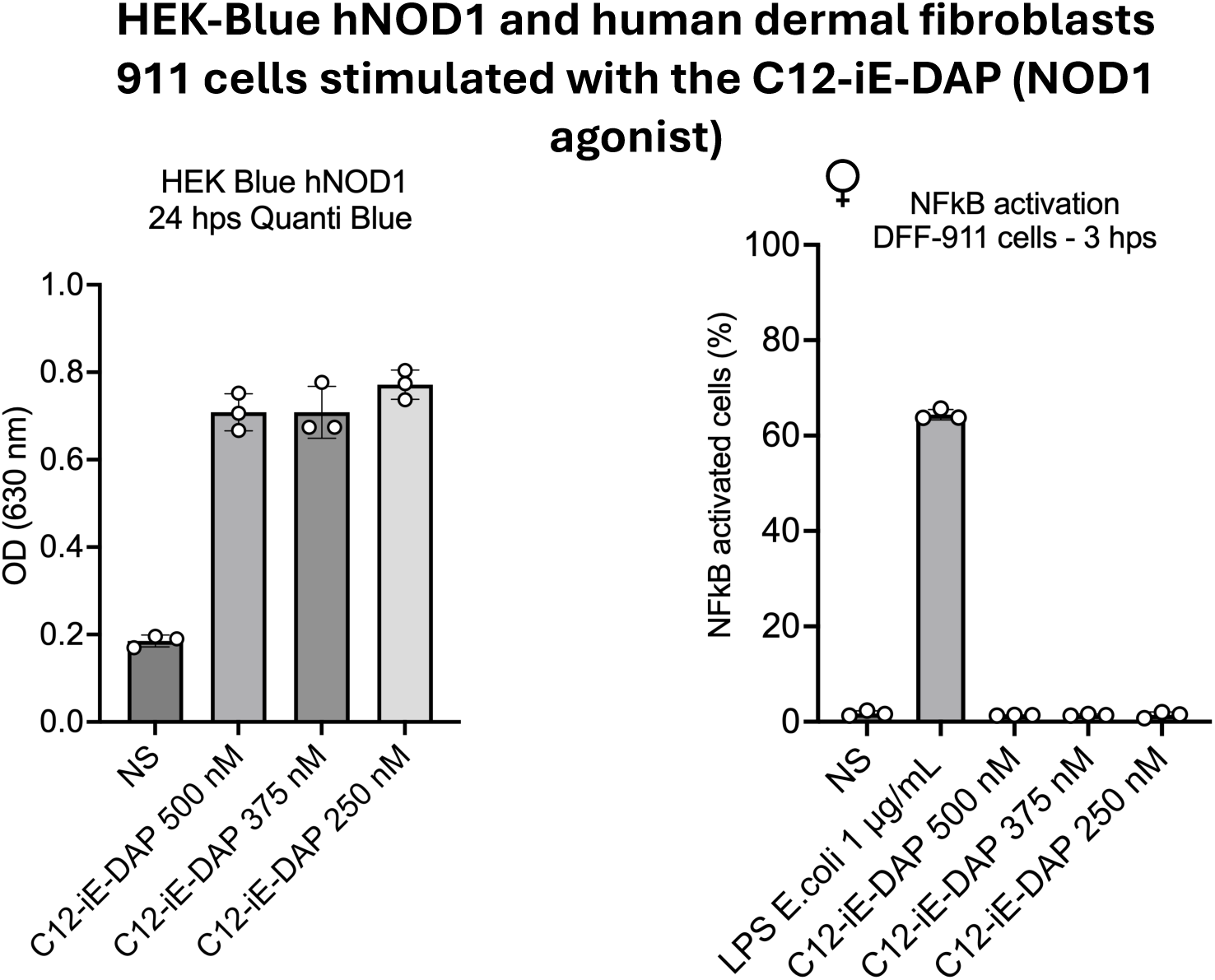
HEK-Blue hNOD1 and dermal fibroblasts (DFF-911) stimulated with C12-iE-DAP. HEK-Blue hNOD1 cells (InvivoGen) and human dermal fibroblasts (DFF-911) were stimulated with different concentrations of the NOD1 agonist C12-iE-DAP (InvivoGen) respectively for 24h and 3h. N=1 with three biological replicates per condition.

**Data sup 5.**
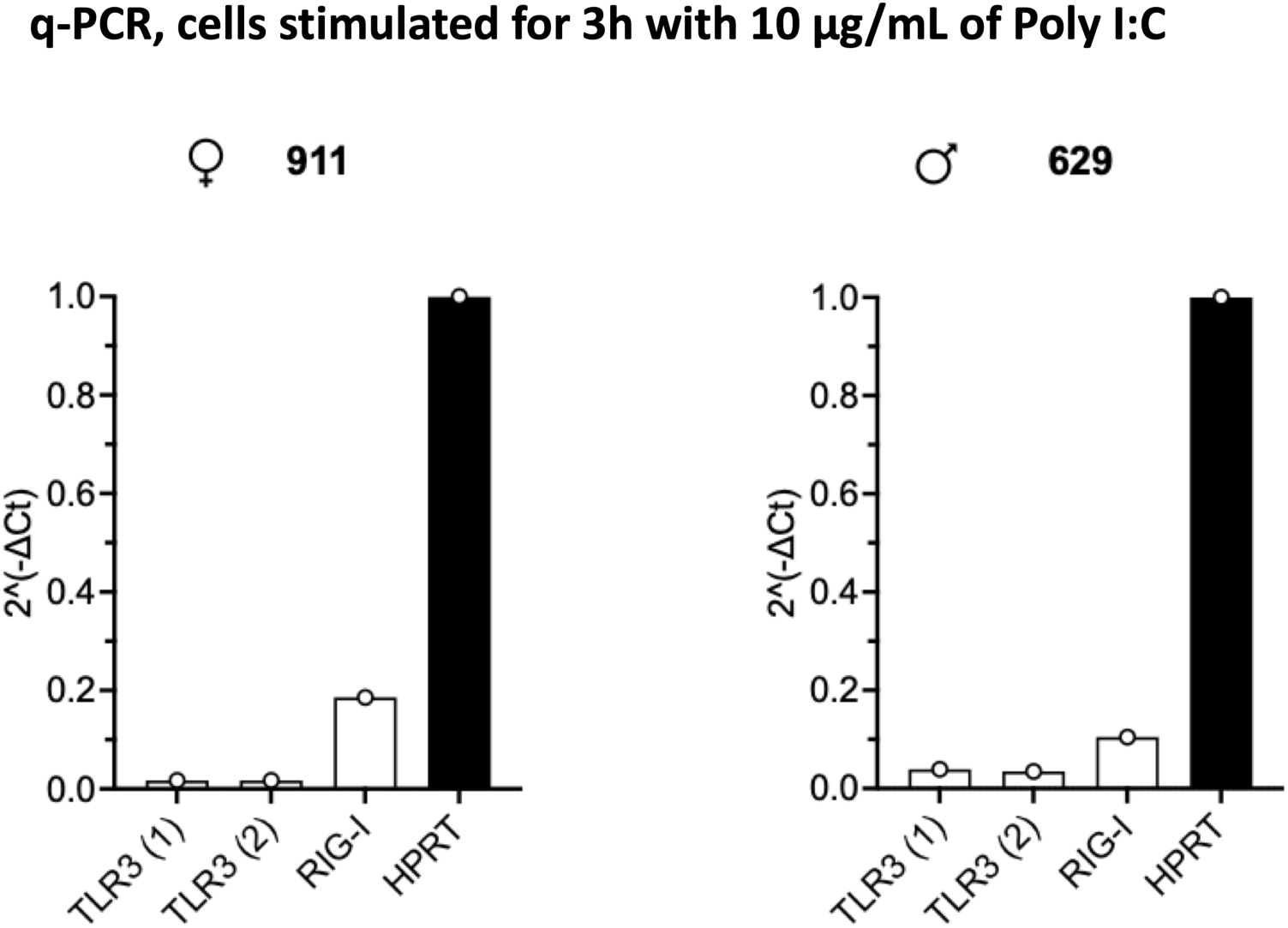
PCR analysis of PRR expression in human dermal fibroblasts. Expression of MDA5, RIG-I and TLR3 in stimulated primary human dermal fibroblasts for 3h with 10 µg/mL of Poly I: C. TLR3 (1) and TLR3 (2) correspond to a different reference of probe (Table 2). mRNA levels were quantified by RT-qPCR in fibroblasts from a female donor (DFF-911) and a male donor (DFF-629). Expression is shown as 2^(-ΔCt) relative to HPRT (set to 1). Each data point represents one fully independent biological replicate, consisting of an independent cell culture, RNA extraction, cDNA synthesis and PCR reaction

## Notes

### Competing Interest Statement

The authors have declared no competing interest.

